# Speed fluctuations of bacterial replisomes

**DOI:** 10.1101/2021.10.15.464478

**Authors:** Deepak Bhat, Samuel Hauf, Charles Plessy, Yohei Yokobayashi, Simone Pigolotti

## Abstract

Replisomes are multi-protein complexes that replicate genomes with remarkable speed and accuracy. Despite their importance, their dynamics is poorly characterized, especially in vivo. In this paper, we present an approach to infer the replisome dynamics from the DNA abundance distribution measured in a growing bacterial population. Our method is sensitive enough to detect subtle variations of the replisome speed along the genome. As an application, we experimentally measured the DNA abundance distribution in *Escherichia coli* populations growing at different temperatures using deep sequencing. We find that the average replisome speed increases nearly five-fold between 17°C and 37°C. Further, we observe wave-like variations of the replisome speed along the genome. These variations correlate with previously observed variations of the mutation rate. We interpret this correlation as a speed–error trade-off in DNA replication. Our approach has the potential to elucidate replication dynamics in *E. coli* mutants and in other bacterial species.

## Introduction

Every cell must copy its genome in order to reproduce. This task is carried out by large protein complexes called replisomes. Each replisome separates the two DNA strands and synthesizes a complementary copy of each of them, thereby forming a Y–shaped DNA junction called a replication fork. The speed and accuracy of replisomes is impressive (Baker and Bell, 1998). They proceed at several hundreds to one thousand base pairs per second (Pham et al., 2013; Elshenawy et al., 2015), with an inaccuracy of about one mis-incorporated monomer every 10 billion base pairs (Schaaper, 1993). In bacteria, two replisomes initiate replication at a well-defined origin site on the circular genome, progress in opposite directions and complete replication upon encountering each other in a terminal region.

The initiation and the completion of DNA replication conventionally delimit the three stages of the bacterial cell cycle (Dewachter et al., 2018; Wang and Levin, 2009). The first stage, B, spans the period from cell birth until initiation of DNA replication. The second stage, C, encompasses the time needed for replication. The last phase, D, begins at the end of DNA replication and concludes with cell division. While it is established that DNA replication and the cell cycle must be coordinated, their precise relation has been a puzzle for decades (Willis and Huang, 2017). A classic study by Cooper and Helmstetter (Cooper and Helmstetter, 1968) finds that, upon modifying the growth rate by changing the nutrient composition in *Escherichia coli*, the durations of stages C and D remain constant at about 40 min and 20 min, respectively. This means that the replisome speed must be unaffected by the nutrient composition, at least on average. When the cell division time is shorter than one hour, DNA replication is initiated in a previous generation. This implies that, in fast growth conditions, multiple pairs of replisomes simultaneously replicate the same genome (Fossum et al., 2007). Tuning the growth rate by changing the temperature has a radically different effect on bacterial physiology. For example, in vivo (Pierucci, 1972) and in vitro (Yao et al., 2009) studies show that the speed of replisomes is affected in this case.

More precise features of replisome dynamics, such as whether their speed is approximately constant or varies along the genome, are important to determine the location of their encounter point and the distribution of replication errors on the genome Niccum et al. (2019); Dillon et al. (2018). However, this detailed information is hard to obtain (Pham et al., 2013). One way for inferring it is to measure the DNA abundance distribution, i.e. the frequency of DNA fragments along the genome in an exponentially growing cell population. In fact, the frequency of these fragments in the population depends on the proportions of synthesizing genomes of different lengths, which in turn depend on the replisome dynamics. Previous studies have measured the DNA abundance distribution, but focused on qualitative analysis of the observed variations in knockout mutants (Wendel et al., 2014, 2018; Rudolph et al., 2013; Midgley-Smith et al., 2018b,a).

In this paper, we introduce a method to infer the replisome dynamics from the DNA abundance distribution. As an application, we experimentally measured the DNA abundance distribution of *E. coli* growing at different temperatures between 17°C and 37°C using high throughput sequencing. Our approach, combined with our experiments, shows that the average speed of replisomes exhibits an Arrhenius dependence on the temperature, with an almost five-fold variation in the range we considered. Moreover, the precision of our experiments reveals that the speed of replisomes varies along the genome in a seemingly periodic and highly repeatable fashion around this average value. We find that this pattern is highly correlated with previously observed wave-like variations of the single base pair mutation rate along the bacterial genome (Niccum et al., 2019; Dillon et al., 2018). We discuss possible common causes for these two patterns.

## Results

### Distribution of genome types

We consider a population of bacteria that grow exponentially in a steady, nutrient-rich environment. Each cell in the growing population can encompass three types of genomes, see Figure 1a and Figure 1b: (i) one template genome, i.e., the genome that the cell inherited at its birth. (ii) incomplete genomes, i.e., genomes which are being synthesized. (iii) post-replication genomes that will be passed to new cells and become their templates.

**Figure 1:**
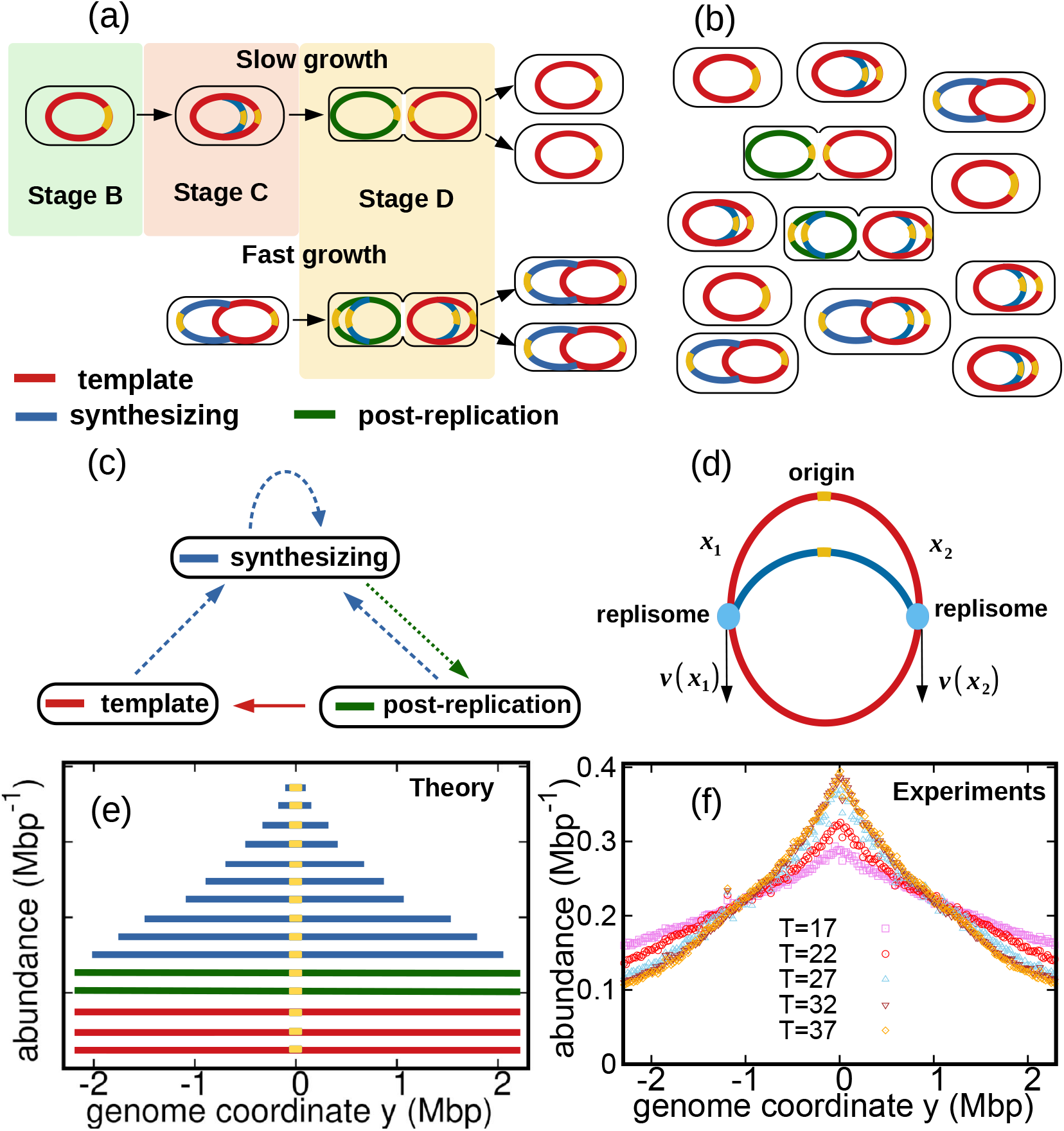
Genome types, their dynamics and DNA abundance distribution in an exponentially growing population. (a) Cell cycle: In slow growth conditions (top panel), newborn cells contain a template (stage B, red). As cell cycle progresses, replisomes begin synthesis of a new genome (stage C, blue) from the origin on the template (yellow spot). When replication terminates, cells contain the original template and a post-replication genome (stage D, green). Upon subsequent cell division, the post replication genome becomes the template for the newborn cell. In fast growth conditions (bottom panel), newborn cells acquire a template which is already undergoing synthesis. (b) Composition of genomes in an exponentially growing population of cells. Each cell may contain a different number of genomes, depending on its stage in the cell cycle and growth conditions. (c) Dynamics of genome types. Dashed blue arrow represent initiation of replication. Dotted green arrow represent completion of replication. Solid red arrow represent cell division. (d) Replisome dynamics. Two replisomes begin replication at an origin and progress in opposite direction. Their speed may depend, in general, on their genome coordinate. (e) Sketch of the DNA abundance distribution as a function of the genome coordinate. All three types of genomes contribute to the DNA abundance distribution. Because of incomplete genomes, the DNA abundance is largest at the origin and smallest at the terminal region (i.e., towards the periphery). (f) Experimental DNA abundance distribution at different temperatures.

In nutrient-rich conditions, bacteria replicate their genome in parallel, so that the numbers of incomplete genomes and post-replication genome per cell are variable, see Figure 1b. The classic Cooper-Helmstetter model (Cooper and Helmstetter, 1968) describes the dynamics of these genomes in a given cell through generations. We adopt a different approach and focus on the abundance dynamics of the three types of genomes in the whole population. We call *N_T_*(*t*), *N_S_*(*t*), *N_P_*(*t*) the total number of template genomes, incomplete (synthesizing) genomes, and post-replication genomes, respectively, that are present in the population as a function of time *t*. Our aim is to quantify the relative fractions of these three types of genomes.

The total number of genomes is *N*(*t*) = *N_T_*(*t*) + *N_S_*(*t*) + *N_P_*(*t*). Since each cell contains exactly one template, the total number of cells is equal to *N_T_*(*t*). The total number of genomes evolve as effect of: (a) replication initiation, which creates a new synthesizing genome at a rate *k*; (b) completion of replication, which transforms a synthesizing genome into a post-replication one at rate *β*; and (c) cell division, which turns a post-replication genome into a template at a rate *α*, see Figure 1c. The model predicts that, in steady growth, the number of cells increases exponentially at a rate equal to the fork firing rate *k* (see Methods).

We now analyze the incomplete genomes in detail. We call *x*_1_ and *x*_2_ the portions of a given incomplete genome copied by the two replisomes at a given time, with 0 ≤ *x*_1_, *x*_2_ ≤ *L*, see Figure 1d. Replication initiates at *x*_1_ = *x*_2_ = 0 and completes once the replisomes meet each other, i.e. *x*_1_ + *x*_2_ = *L*.

The replisome dynamics proceeds as follows. Each replisome is characterized by a speed *v*(*x*) which depends, in principle, on the replisome position *x* (be it *x*_1_ or *x*_2_) and by a diffusion coefficient *D* representing random fluctuations of the speed, see Methods. Close to thermodynamic equilibrium, the diffusion coefficient D can be estimated by the Stokes-Einstein relation (Hynes, 1977). However, since replisomes are driven far from equilibrium by the hydrolysis of dNTPs, their diffusion coefficient could deviate from this estimate. In the absence of fluctuations (*D* = 0), each of the two replisomes copies exactly half of the genome, whereas for *D* > 0 their meeting point is characterized by a certain degree of uncertainty.

In steady exponential growth, the copied portions *x*_1_, *x*_2_ of incomplete genomes are characterized by a stationary probability distribution *p*^st^(*x*_1_, *x*_2_), which depends on the replisome dynamics.

### DNA abundance distribution

The DNA abundance distribution 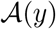 is the probability that a small DNA fragment randomly picked from the population originates from genome position *y*, see Figure 1c. We define the genome coordinate *y* ∈ [–*L*/2,*L*/2], where *y* = 0 corresponds to the origin of replication and *L* is the genome length. Templates and post-replication genomes yield a uniform contribution to the distribution 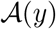 (red and green in Figure 1c). In contrast, incomplete genomes contribute in a way that depends on the replisome positions along the genomes (blue in Figure 1c). Our experiments permit to measure the distribution 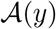 with high accuracy, see Figure 1d and Methods.

For given choices of *v*(*x*) and *D*, our theory permits to compute the distribution of incomplete genomes *p*^st^(*x*_1_, *x*_2_) and thereby the DNA abundance distribution 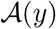, see Methods. Therefore, by experimentally measuring the DNA abundance distribution we can test our hypotheses on the speed function *v*(*x*) and the diffusion coefficient *D*.

### Constant speed model

We first consider a scenario in which replisomes progress at a constant speed 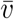 and without fluctuations, *D* = 0. We find that, in this case, the DNA abundance distribution is expressed by

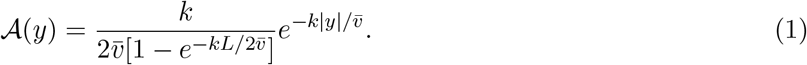

We fit this distribution to the experimental data using maximum likelihood, see Figure 2a. The speed *v* is the only fitting parameter as we independently measure the exponential growth rate *k* from the optical density, see Supporting Figure S1.

**Figure 2:**
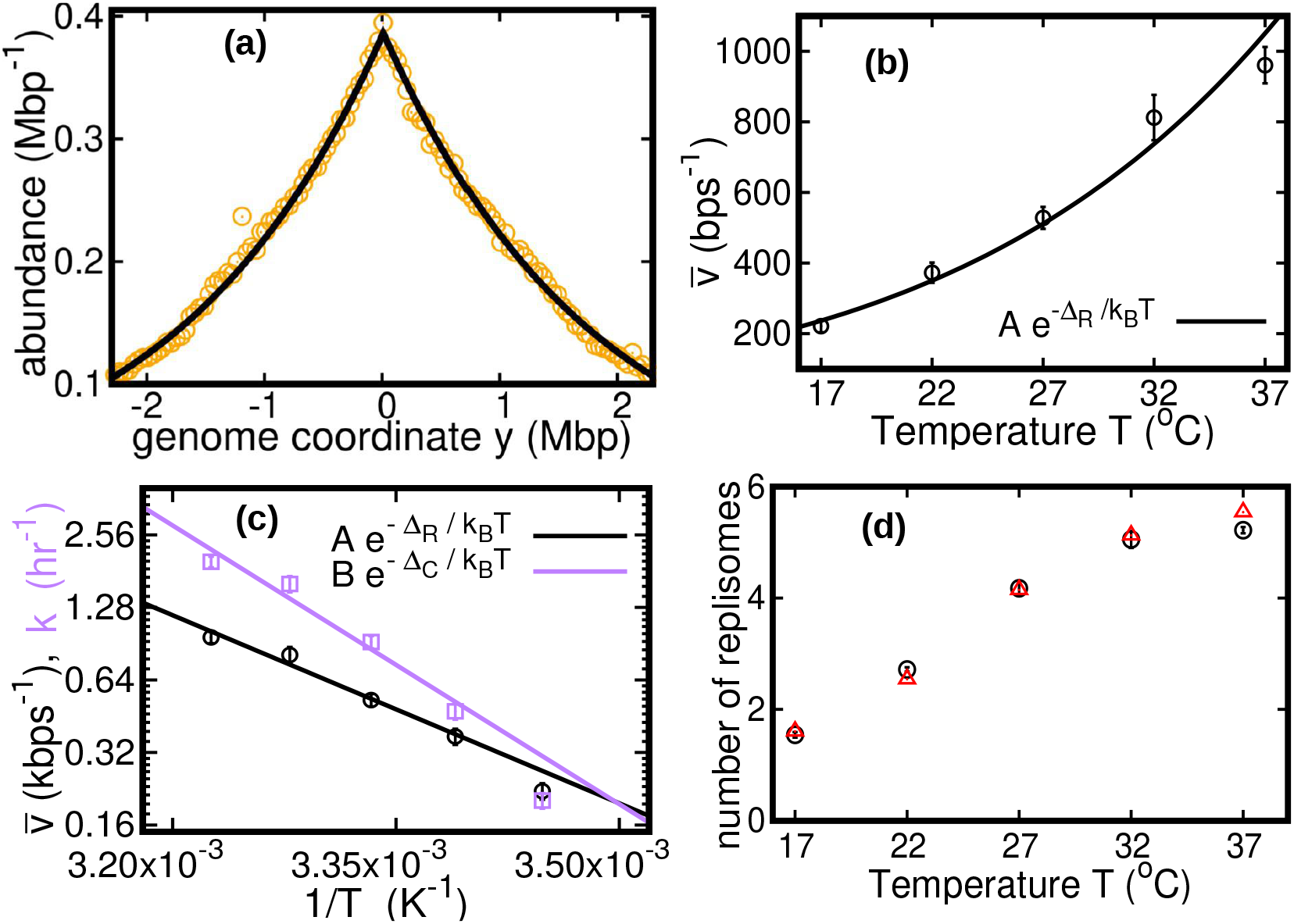
Results of the constant speed model. (a) DNA abundance distribution for *T* = 37°C. Orange circles represent experimental data. The solid black line is the prediction of our model assuming constant speed and *D* = 0. Fits are performed using a maximum likelihood method, see Appendix A for details. The quality of fits for replicates and other temperatures is comparable, see Supporting Figure S2, Supporting Figure S3 and Supporting Figure S4. In particular, fits of replicates yield similar values of the speed 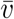. (b) Replisome speed as a function of temperature. Error bars represent sample-to-sample variations. (c) Comparison of the temperature-dependence of speed and growth rate (see Supporting Figure S1). The solid curves are fits of Arrhenius laws to the data. The fitted parameters are *A* = (2.5 ± 5.3) × 10^8^bps^−1^, Δ_*R*_ = (50 ± 5)kJmol^−1^, *B* = (6.0 ± 24.9) × 10^12^hr^−1^ and Δ_*C*_ = (74 ± 10)kJmol^−1^. We exclude the data point for *T* = 17°C in both fits. (d) Estimated number of replisomes per complete genome at different temperature. The red triangles represents the estimate from eq. (17) in which we use the expression of *β* for the constant speed model, eq. (20). The black circles are the estimates from eq. (18).

We find that the speed increases nearly five fold with temperature in the range we considered, see Figure 2b. This temperature dependence appears to follow an Arrhenius law, see Figure 2b. This behavior resembles that of the growth rate. The effective activation energy characterizing the cell cycle is larger than that characterizing the replisome speed, see Figure 2c, possibly due the large number of molecular reactions involved in the cell cycle. The data point at 17°C appears to deviate from the Arrhenius law for both the speed and the growth rate (Roy et al., 2021), see Figure 2c.

In fast growth conditions (i.e., high temperature), multiple replisomes synthesize DNA in parallel inside each cell. In the temperature range we studied, we predict that the number of replisomes per complete genome increases from two to almost six, see Figure 2d.

The average DNA content per cell 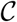 is the product of the average genome length ℓ times the average number of genomes per cell *N*/*N_T_*. Using eq. (16) for the average genome length and eq. (9) for the average number of genomes per cell, we obtain that the average DNA content per cell is expressed by

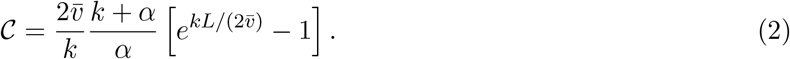

Since we assumed constant speed and *D* = 0, the duration of the replication cycle is equal to 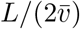 and does not fluctuate. The classic Cooper-Helmstetter model (Cooper and Helmstetter, 1968) predicts the DNA content per cell assuming constant durations of stages B, C, and D of the cell cycle. Indeed, if we make the same assumptions, we find that the prediction of eq. (2) becomes equivalent to that of the Cooper-Helmstetter model (see Appendix B).

The average DNA content per cell gives us a chance to discuss the notion that the ratio between the amount of DNA and the amount of proteins per cell should be maintained approximately constant at varying the growth rate (Si et al., 2017). If the growth rate is varied by changing the nutrient composition, 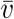 remains constant, and eq. (2) predicts an approximately exponential growth of 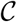 with *k*, which is consistent with observations (Cooper and Helmstetter, 1968). In this case, the Schaechter–Maaloe–Kjeldgaard growth law states that the cell size grows exponentially with *k* as well (Schaechter et al., 1958), thereby ensuring DNA-protein homeostasis. In the case of varying temperature, we find that 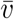 and *k* present a similar dependence on *T* (see Figure 2c), so that their ratio and thereby 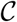 weakly depends on *k* (see Appendix B Figure B1). Our result is consistent with the observation that, at varying temperature, the cell size remains approximately constant as well (Shehata and Marr, 1975; Trueba et al., 1982).

### Oscillating speed model

The assumption of constant speed leads to a rather good fit of our DNA abundance data. However, the precision of our sequencing data permits us to appreciate systematic deviations from the model predictions under the constant speed hypothesis, see Figure 3(a-e). These deviations appear as regular oscillations as a function of the genome coordinate. They are highly repeatable (see Supporting Figure S5, Supporting Figure S6 and Supporting Figure S7) and approximately symmetric with respect to the origin of replication. Our analysis reveals similar oscillations in experimental data from a previous study (Midgley-Smith et al., 2018b), see Supporting Figure S8. Taken together, these evidences support that this phenomenon is very robust.

**Figure 3:**
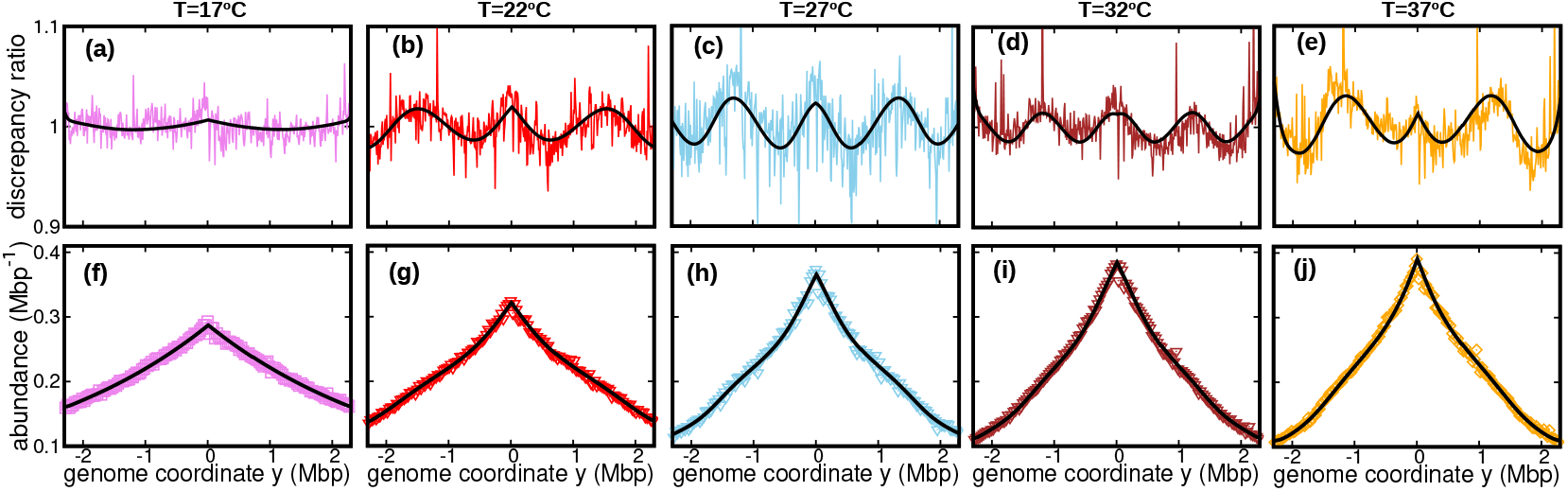
Deviations from the predictions of the constant speed model indicate that the replisome speed oscillates along the genome. (a-e) Colored lines: ratios of the experimental DNA abundance over the corresponding prediction assuming constant speed and *D* = 0. These ratios exhibit a wave-like pattern. The solid black lines represent the ratios of the predictions assuming oscillatory speed, eq. (3) and *D* ≥ 0, over constant speed and *D* = 0. Corresponding plots for replicates and other temperatures are presented in Supporting Figure S5, Supporting Figure S6 and Supporting Figure S7. (f-j) Experimental DNA abundance distribution at different temperatures. The solid black lines are the fits of the oscillatory speed model. Tests based on the Akaike information criterion show that the oscillatory speed model should be chosen over the constant speed model for all the replicates at different temperatures, see Supporting Figure S9. The fitted parameters are reported in Table 1.

To account for this observation, we introduce a more refined model in which the replisome speed oscillates along the genome:

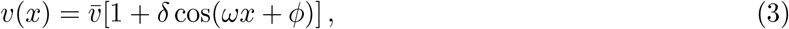

where *δ* represent the relative amplitude of oscillations; *ω* their angular frequency along the genome; and *ϕ* their initial phase. We also take into account random speed fluctuations in this case, *D* ≥ 0. By fitting the DNA abundance, we estimate the parameters 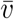, *δ, ω*, *ϕ*, and *D*, see Figure 3(f-j) and Table 1.

**Table 1:**
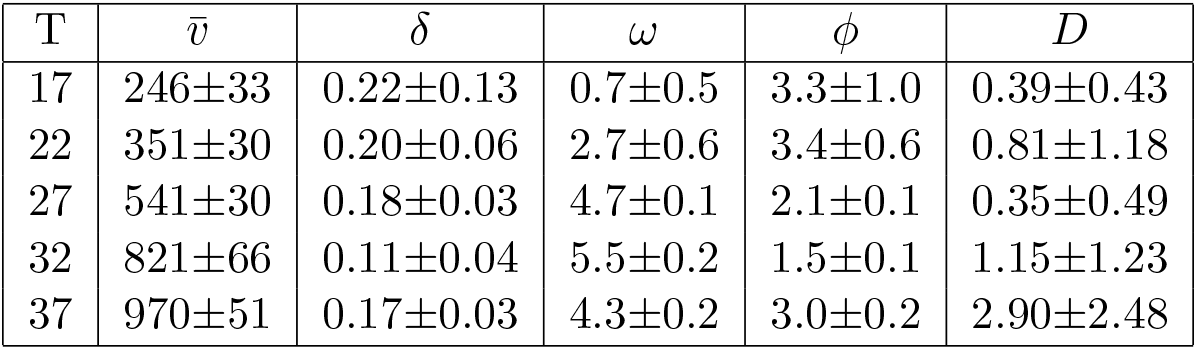
Parameters of the oscillatory speed model. Temperatures are expressed in °C, 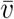 in bps^−1^, *ω* in rad Mbp^−1^, *ϕ* in radians and *D* in kbp^2^s^−1^. Reported values are average over experimental replicates. Error bars represent sample-to-sample variations. The average speed estimated from the oscillatory speed model and constant speed model are comparable, see Appendix C Table C1.

Our fitted speed oscillations are reminiscent of a previously observed wave-like pattern in the mutation rate along the genome of different bacterial species (Dillon et al., 2018; Niccum et al., 2019). For a quantitative comparison, we analyze this pattern in a mutant *E. coli* strain lacking DNA mismatch repair (Niccum et al., 2019). We find that the oscillations in mutation rate and in speed are highly correlated, see Figure 4a. The mutation rate appears approximately in phase with the speed, meaning that regions where replisomes proceed at higher speed are characterized by a higher mutation rate. This observation strongly suggests that the two phenomena have a common cause.

**Figure 4:**
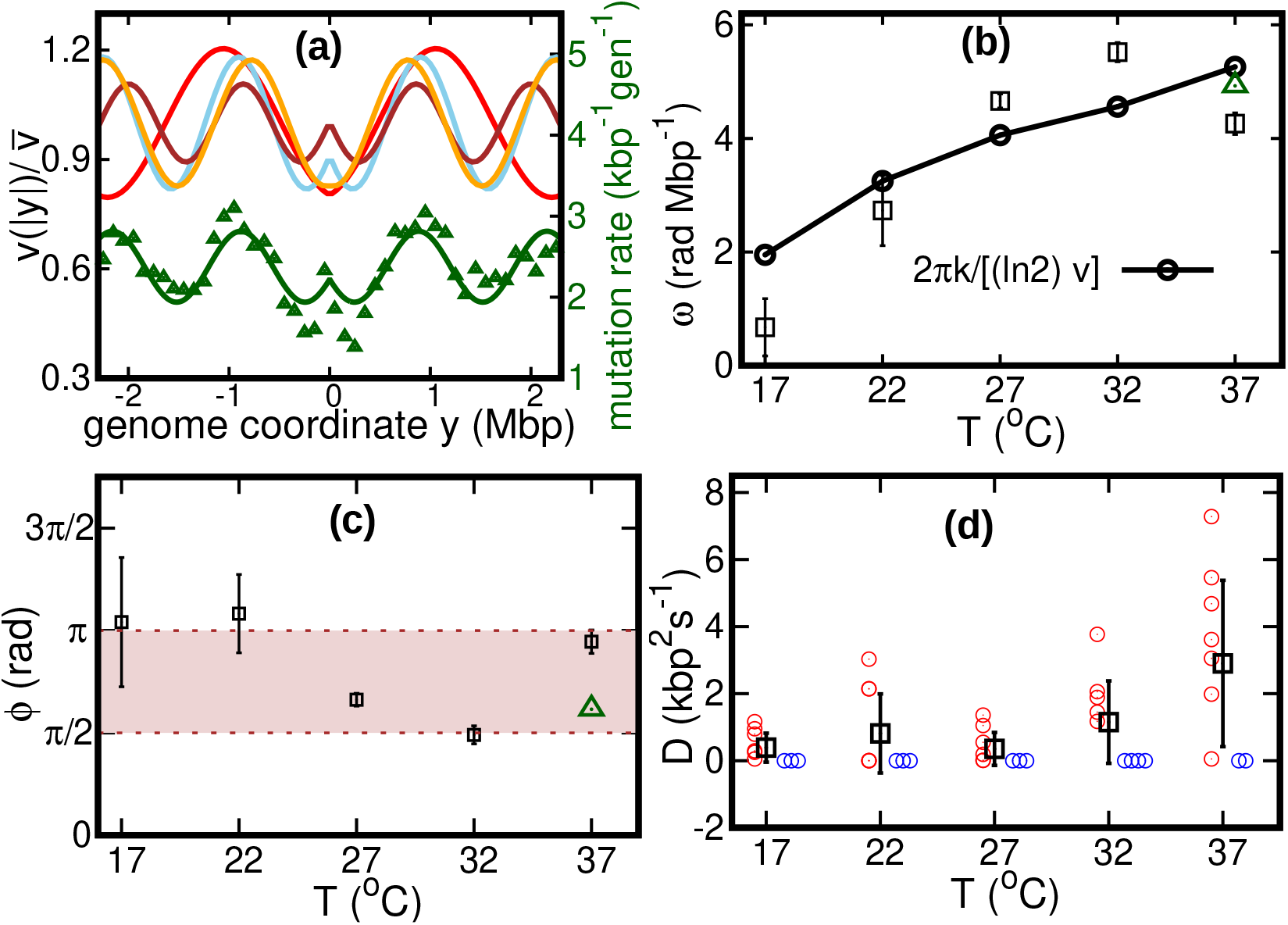
Results of the oscillatory speed model. (a) Solid lines: relative speeds 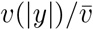 along the genome (Red: *T* = 22°C, sky blue: *T* = 27°C, brown: *T* = 32°C, and orange: *T* = 37°C). We omitted the curve for *T* = 17°C as the oscillations are less evident in this case (see Supporting Figure S10). The wave-like pattern of the speed is quantitatively similar to the variations of the mutation rate along the genome (green triangles, from (Niccum et al., 2019); Pearson correlation coefficients between speed and mutation rate: *r*_22*C*_ = 0.42; *r*_27*C*_ = 0.84; *r*_32*C*_ = 0.80; and *r_37*C*_* = 0.69). The mutation rate is defined as the number of base pair substitutions per generation per kilo base pairs. The solid green line is a fit to the mutation rate data with a function that has same form as in (3). The fit parameters are 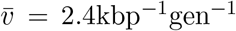, *δ* = 0.18, *ω* = 4.9radMbp^−1^ and *ϕ* = 1.93rad. (b) Temperature dependence of angular frequency of oscillation *ω*. (c) phase *ϕ*. Green triangles in (b) and (c) represent the angular frequency and phase, respectively, from the fit to the mutation rate data with eq. (3). (d) Diffusion coefficient *D*. Circles represent individual fitted values of diffusion coefficients. Blue circles represent cases in which the fitted value of *D* is either zero or not significant (see SI). This occurs in 2 out of 9 cases for 37°C and 3 out of 9 cases for the other temperatures.

We consider two possible explanations for these oscillations. The first is that the oscillations originate from a systematic process related with the cell cycle (Niccum et al., 2019). The second explanation is that the oscillations are caused by competition among replisomes for nucleotides or other molecules required for replication. Assuming approximately constant cell division times, we estimate the cell division time as *τ* = (ln 2)/*k*. Since *k* is also equal to the fork firing rate per genome, the time between firing events in a cell is also approximately equal to *τ*, so that the two hypotheses lead to the same quantitative prediction. If the speed of replisomes was coupled to a factor oscillating with period *τ*, this would cause spatial oscillation of speed with angular frequency 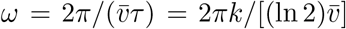. This prediction qualitatively agrees with our fitted values of *ω*, see Figure 4b.

If the wave-like pattern were caused by competition among replisomes, one would expect either a minimum of the speed every time a new fork is fired (*ϕ* = π) or the speed to start decreasing when a new fork is fired (*ϕ* = π/2). Our fitted values of the phase *ϕ* are compatible with this range, see Figure 4c.

Our results show that the diffusion coefficient *D* is quite small. For about one third of our experimental realizations at each temperature, our fitted value of *D* is not significant according to the Akaike information criterion (see Figure 4d and Supporting Figure S11). For comparison, we estimate the equilibrium diffusion constant of replisomes in the cytoplasm from the Stokes-Einstein relation as *D*_SE_ ≈ 6kbp^2^s^−1^ (see Appendix D), of the same order of magnitude of our fitted values, see Table 1 and Figure 4. Taken together, these results suggest that, despite their high speed, the fluctuations of replisome position are remarkably similar to the equilibrium case.

The diffusion coefficient determines the uncertainty about the genome site where the two replisomes meet. In the absence of diffusion (*D* = 0), replisomes would always meet at the mid point on the circular genome. For *D* > 0, we estimate the typical size *l*_D_ of the region in which the two replisomes meet as follows. Since the fitted values of *δ* and *D* are both small, we approximate the replication time as 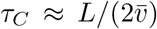. In this time, the accumulated uncertainty due to diffusion is equal to 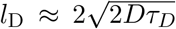. From our estimated diffusion coefficients and average velocities, we obtain values of *l*_D_ on the order of 100 – 200kbp, depending on temperature.

## Discussion

In this paper, we developed a theory to infer the dynamics of replisomes from the DNA abundance distribution in a growing bacterial population. We tested our method by measuring the DNA abundance distribution of growing *E. coli* populations at different temperatures. We found that the dependence of the average speed on the temperature is well described by an Arrhenius law, similar to that governing the population growth rate. Our analysis shows that this result is compatible with DNA-protein homeostasis, given that the average cell size hardly varies with temperature (Shehata and Marr, 1975; Trueba et al., 1982).

Our approach reveals a wave-like oscillation of the replisome speed along the *E. coli* genome. The relative amplitude of these oscillations ranges from 10% to 20% of the average replisome speed. A quantitatively similar pattern was observed in the bacterial mutation rate along the DNA of an *E. coli* mutant strain (Niccum et al., 2019) and in other bacterial species (Dillon et al., 2018). This similarity strongly suggests that the two phenomena have a common dynamical origin. In particular, we suggest that their link could be the trade-off between accuracy and speed that characterizes DNA polymerases (Sartori and Pigolotti, 2013; Banerjee et al., 2017; Fitzsimmons et al., 2018). Because of this tradeoff, any mechanism increasing the speed of a polymerase is expected to reduce its accuracy as well. Our analysis of the wave length of these oscillations supports that this pattern originates from a process synchronized with the cell cycle (Dillon et al., 2018), whose activity alters the replisome function. An alternative explanation is that the oscillations originate from competition among replisomes for shared resources.

Beside these regular and repeatable fluctuations, our analysis shows that random fluctuations of replisome speed are quite small, leading to an uncertainty of about 100 – 200k*bp* on the location of the replisome meeting point. In bacteria, the terminal region of replication is flanked by two groups of termination (Ter) sites having opposite orientation. Ter sites are binding site for the Tus protein and permit passage of replication forks in one direction only (Elshenawy et al., 2015), so that the two groups effectively trap the two forks in the terminal region (Duggin et al., 2008). Out of the ten Ter sequences in *E. coli*, only two of them (TerB and TerC) are within 100 – 200k*bp* of the point diametrically opposite to the origin. These two sequences have the same orientation. Our result therefore implies that most Ter sequences are usually not needed to localize the replisome meeting point. This prediction is consistent with previous observations that the phenotypes of Tus- *E. coli* mutants (Roecklein et al., 1991) or mutants lacking Ter sequences (Duggin et al., 2008) do not appear distinct from that of the wild type.

Quantitative modeling of the DNA abundance distribution has the potential to shed light on aspects of replisome dynamics beyond those explored in this paper. For example, it was observed that knockout of proteins involved in completion of DNA replication leads to either over-expression or under-expression of DNA in the terminal region (Wendel et al., 2014, 2018; Sinha et al., 2018). Incorporating the role of these proteins into our model will permit to validate possible explanations for these patterns. More in general, our approach is simple and general enough to be readily applied to other bacterial species, to unravel common principles and differences in their DNA replication dynamics.

## Methods

### Cultivation and DNA extraction

*E. coli* MG1655 was cultured in LB medium supplemented with 50mM MOPS pH 7.2 and 0.2% glucose. Overnight cultures grown at 37°C were diluted into fresh medium and grown until reaching an OD600 of about 1.0 at the target temperature. These cultures were used to inoculate 50 ml medium at the desired temperature in 500ml Erlenmeyer flasks with baffles at a target OD of 0.01. Cultivation was performed with shaking at 250 rpm. OD was determined with a NanoDrop One in cuvette mode.

*E. coli* cultures (1.4 ml) were harvested by centrifugation at 21000g for 20 seconds after reaching an OD of around 0.5 (mid-exponential phase, see Supporting Figure S1). Cells were kept growing for at least 45 doubling times (as measured in exponential phase) to reach stationary phase. Samples of 0.2 ml from the stationary phase cultures grown at 17°C, 27°C, and 37°C were harvested for DNA extraction. The pellets were immediately frozen at −80°C until DNA extraction. DNA was extracted in parallel using Genomic DNA Purification Kit from Thermo Fisher Scientific.

### Sequencing

We sequenced three samples in the exponential phase from different experimental realizations for each temperature. In addition, we sequenced three stationary samples at three different temperatures. DNA samples were sheared by ultrasound using Covaris AFA technology. Libraries were prepared using the Illumina DNA PCR-Free Library Prep Kit. Sequencing was performed on a Novaseq6000 using paired-end 150 bp reads.

### Alignment and bias elimination

We aligned reads from each sample using Bowtie2 2.3.4.1 (Langmead and Salzberg, 2012), using the MG1655 genome as a reference. We calculated the frequency of reads as a function of the genome coordinate with bin size 10k*bp*. To attenuate bias, we divided the frequency at each genome coordinate in a sample from the exponential phase by the frequency of the corresponding bin in a stationary sample (Wendel et al., 2014; Midgley-Smith et al., 2018b). We alternatively used all of our three stationary samples to correct the bias of each sample in the exponential phase. Therefore, after bias elimination, we effectively have 3 × 3 = 9 different DNA abundance curves in the exponential phase at each temperature. See Supporting Figure S2, Supporting Figure S3 and Supporting Figure S4 for details.

### Distribution of genome types

In this subsection we compute the relative fractions of the different genome types. The genome numbers evolve as

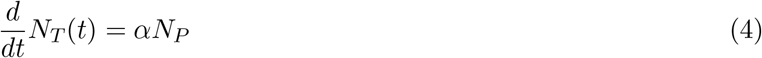

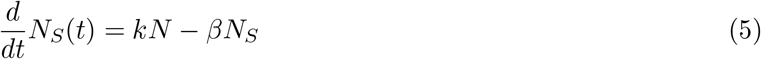

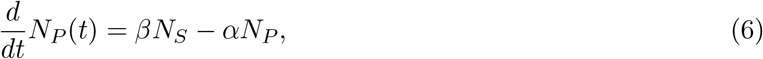

where *α* is the cell division rate per post-replication genome, *k* is the rate per genome at which forks are fired in the population, and *β* is the rate of completion of DNA replication per incomplete genome, see Figure 1c. It follows from eq. (4) that, in steady growth, the total number of genomes grows exponentially at rate *k*, *N*(*t*) ∝ exp(*kt*). In this exponential regime, the fractions of the three genome types are constant:

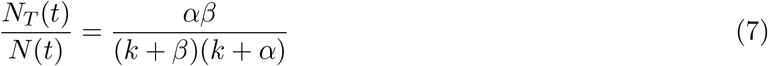

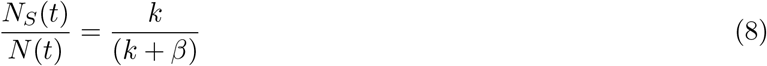

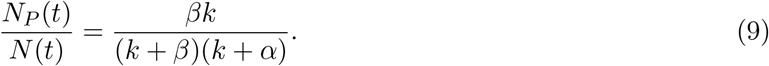

he ratio *N*/*N_T_* can be interpreted as the average number of genomes per cell. Since this ratio is constant, the fork firing rate *k* can also be identified as the exponential growth rate of the number of cells.

### Replisome dynamics

We call *n_S_*(*x*_1_, *x*_2_; *t*) the number density of incomplete genomes at time *t* with replisome positions at *x*_1_ and *x*_2_. By definition 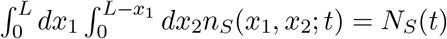. We assume that this number density evolves according to

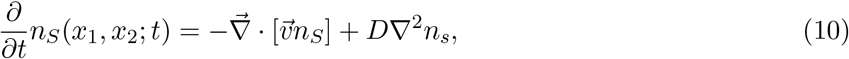

where 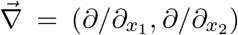 and 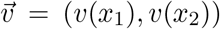. We now introduce the normalized probability *p*(*x*_1_, *x*_2_; *t*) = *n_S_*(*x*_1_, *x*_2_; *t*)/*N_S_*(*t*). By substituting this definition into (10), we obtain

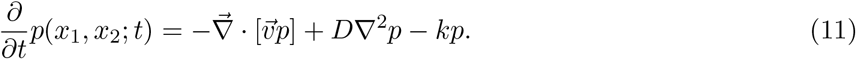

The last term in (11) is a dilution term that accounts for the exponentially increase in newborn cells. The stationary distribution *p*^st^(*x*_1_, *x*_2_) is a time-independent solution of eq. (11).

Because of replication completion, the line *x*_1_ + *x*_2_ = *L* is an absorbing state for the dynamics described by eq. (11). Equation (11) must be consistent with eq. (5), which describes the dynamics of incomplete genomes regardless of the coordinates of their replisomes. This implies that the rate *β* at which replication completes must equal to the probability flux through the absorbing boundary:

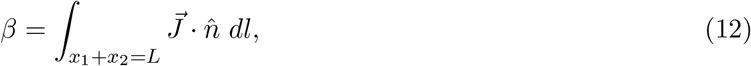

where we introduce the probability current 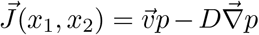, the unit vector 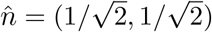, and the infinitesimal line increment *dl* along the absorbing boundary. Similarly, the probability flux entering the system at (*x*_1_, *x*_2_) = (0, 0) must match the rate of replication initiation as given by (5).

### DNA abundance from the replisome distribution

In this section we link the DNA abundance distribution 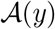 with the replisome position distribution *p*^st^(*x*_1_, *x*_2_). We first introduce the probability 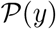 that a randomly chosen genome (either complete or incomplete) in the population includes the genome location *y*. The integral of 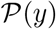 is equal to the average genome length ℓ in the population

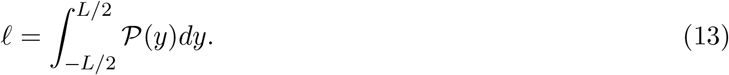

It follows from eq. (8) that a randomly chosen genome is complete with probability (1‒*N_S_*/*N*) = *β*/(*k*+*β*). For incomplete genomes, we take into account that either of the replisomes can have copied position *y*. This argument leads to the expression

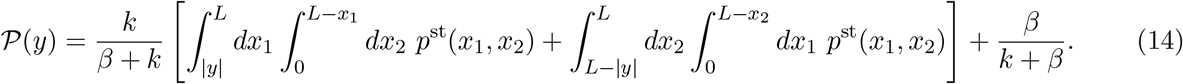

The DNA abundance distribution 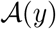 is proportional to 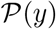, up to a normalization constant:

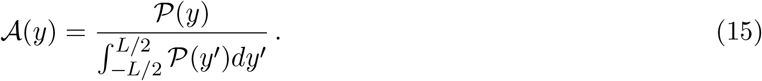

Combining eq. (13), eq. (15), and the fact that 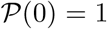, we obtain a simple relation between the DNA abundance distribution and the average genome length:

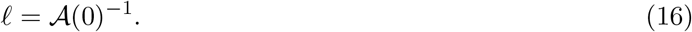

In summary, given an hypothesis on the velocity function *v*(*x*) and the diffusion coefficient *D*, we solve eq. (11) at stationarity using the experimentally measured growth rate *k*. From the stationary solution *p*^st^(*x*_1_, *x*_2_), we obtain *β* using eq. (12). Our approach does not permit to determine the rate *α*. However, this rate is not necessary to compute the DNA abundance distribution, which is expressed by eqs. (14) and (15) in terms of *p*^st^(*x*_1_, *x*_2_) and *β* only.

### Average number of replisomes per complete genome

We estimate the average number of replisomes per complete genome 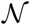 in two alternative ways. On the one hand, using eqs. (7)–(9) we find that

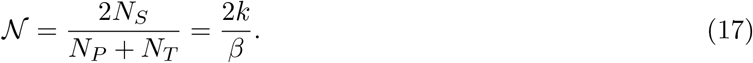

On the other hand, it can be seen in Figure 2d that the fraction of complete genome in the population is equal to the ratio 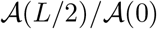 between the DNA abundance at the terminal and at the origin. It follows that

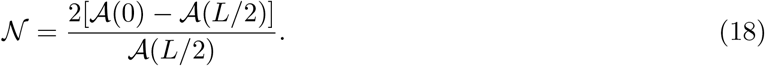

### Constant speed

We focus on the scenario with constant speed and *D* = 0. In this case, the steady solution of eq. (11) is given by

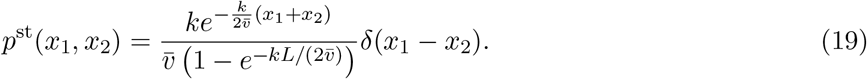

The rate at which replication completes is equal to

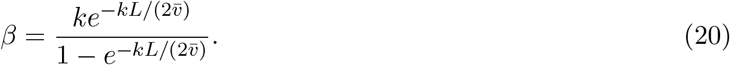

Substituting eqs. (19) and (20) into eq. (14), we obtain

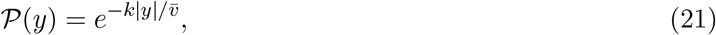

from which eq. (1) follows by normalizing, see eq. (15).

We exactly solved eq. (11) also in the case where the speed oscillates but the diffusion coefficient vanishes, see Appendix E.

### Stochastic simulations

In the case of oscillating speed and *D* > 0, we compute the stationary solution of eq. (11) using numerical simulations. To this aim, we interpret eq. (11) as describing a stochastic process subject to stochastic resetting (Evans and Majumdar, 2011). Specifically, we evolve trajectories according to a system of stochastic differential equations:

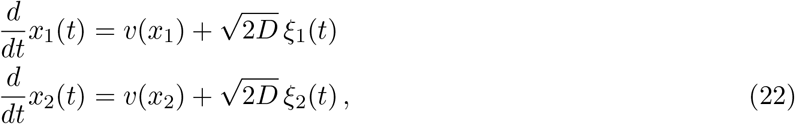

where *ξ*_1_(*t*) and *ξ*_2_(*t*) are white noise sources satisfying 〈*ξ*_1_(*t*)〉 = 〈*ξ*_2_(*t*)〉 = 0, 〈*ξ*_1_(*t*)*ξ*_1_(*t*′)〉 = 〈*ξ*_2_(*t*)*ξ*_2_(*t*′)〉 = *δ*(*t* – *t*′), and 〈*ξ*_1_(*t*)*ξ*_2_(*t*′)〉 = 0. In addition to the dynamics described by eqs. (22), with a stochastic rate *k*, trajectories are reset to the origin, *x*_1_ = *x*_2_ = 0 (blue trajectory in Figure 5a). Since the boundary *x*_1_ + *x*_2_ = *L* is an absorbing state, trajectories that reach this boundary are also reset to the origin (green trajectory in Figure 5a). The probability distribution associated with this dynamics evolves according to eq. (11). We simulate this stochastic dynamics to estimate the stationary distribution *p*^st^(*x*_1_, *x*_2_) in a computationally efficient way, see Figure 5b. We also estimate from the same simulations the parameter *β* as the empirical rate at which the absorbing boundary is reached, see eq. (12).

**Figure 5:**
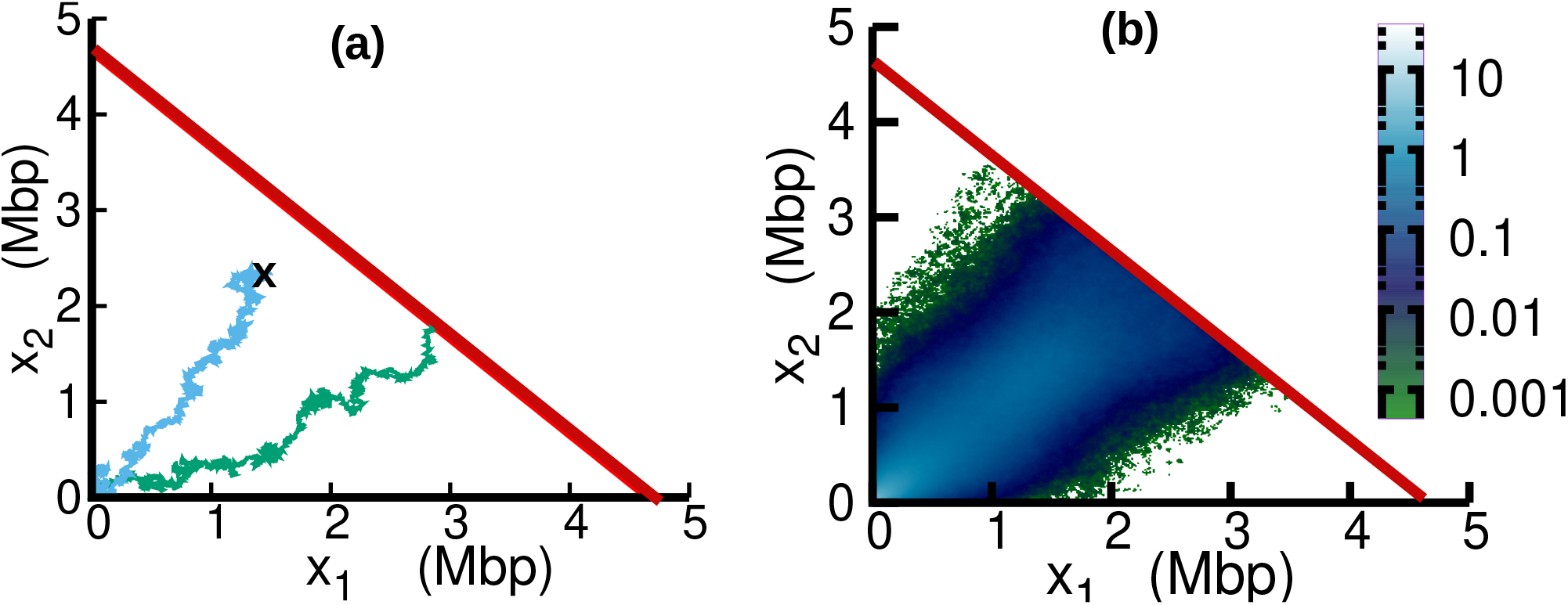
Replisome dynamics in the (*x*_1_, *x*_2_) plane. (a) Two different trajectories demonstrate two different types of resetting events in our simulations. The trajectories are reset to *x*_1_ = 0, *x*_2_ = 0 when either the two replisomes meet (green trajectory) at the absorbing boundary (solid red line) or when a new initiation occurs (sky blue). (b) Replisome position distribution *p*^st^(*x*_1_, *x*_2_) in the steady state. In both panels, parameters are 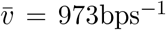, *δ* = 0.19, *ω* = 4radMbp^−1^, *ϕ* = 3.1 and *D* = 55kbp^2^s^−1^. These parameters are on the order of those fitted from experiments (see Table 1), except for *D* which is chosen to be larger for illustration purposes.

## Acknowledgements

We are grateful for the help and support provided by the Scientific Computing and Data Analysis section of Research Support Division at OIST. We thank the DNA Sequencing Section of OIST for support with sequencing. We thank M. Cencini for feedback on a preliminary version of our manuscript.

## Contributions

DB and SP conceptualized the study, developed the model, and carried out the mathematical analysis. DB wrote the computer code and performed the numerical simulations. DB, SH, and YY performed the experiments. CP analyzed the sequencing data. DB and SP wrote the manuscript, with input from SH, CP, and YY.

## Data availability

Sequence reads were deposited in the NCBI Sequence Read Archive with links to BioProject accession number PRJNA772106. Read frequencies on the genome are available in a file archive deposited in Zenodo (DOI:10.5281/zenodo.5577986).

## Appendix A Parameter estimation based on maximum likelihood

In this Section, we outline the the maximum likelihood method to fit the parameters of the model. We make a histogram of our data with bin size *b* = 10kbp. We call *B* the total number of bins. We empirically estimate the DNA abundance in each bin *i* as

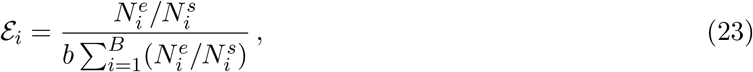

where 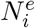 and 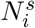 are the numbers of reads in the *i*th bin from the exponential and stationary culture, respectively. We assume that the number of reads in each bin follow a Poisson distribution (Aird et al., 2011). In the limit of large 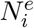 and 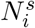, the distribution of the empirical DNA abundance 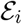 is Gaussian with standard deviation

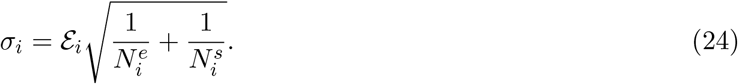

The assumption of large number of reads is well satisfied for our sequencing depth: 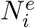 ranges from 2.8 × 10^4^ to 19.9 × 10^4^ and 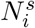 from 4.6 × 10^4^ to 9.0 × 10^4^. We call 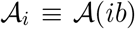 the DNA abundance predicted by our model at bin *i* for a given set of parameters. The joint likelihood of the empirical DNA abundance in all the bins is given by

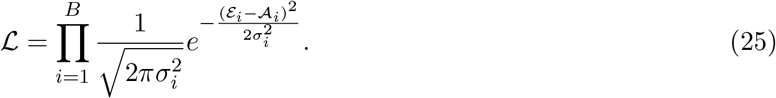

We fit the model parameters by maximizing the logarithm of the likelihood ln 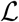.

In the constant velocity case, there is single fitting parameter 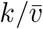. In the oscillatory model without and with diffusion we have four and five fitting parameters, respectively.

## Appendix B Correspondence with the Cooper-Helmstetter Theory

In this Section, we prove the correspondence between our constant velocity model and the Cooper-Helmstetter model in the particular case where the durations of the B and D stages are constant. We additionally assume slow growth conditions, so that all stages B, C, and D of the cell cycle are present.

In particular, we aim to prove that the expression for the total DNA abundance in the constant velocity model

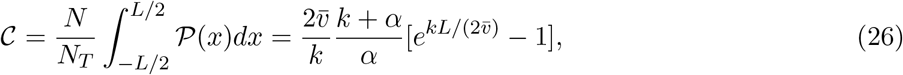

see eq. (2) in the Main Text, is equivalent to the expression for the Cooper-Helmstetter model

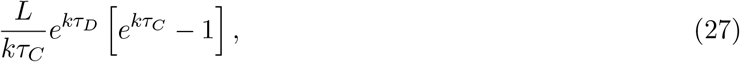

where *τ_C_* is the time taken by replisomes to complete replication on the genome and *τ_D_* is post replication period (Cooper and Helmstetter, 1968). Our proof provides an example of how we can connect population-level parameters such as *α* and *k* to individual-level parameters such as *τ_C_* and *τ_D_*.

Since we assumed constant replisome speed, we have 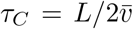. Therefore, proving that eq. (26) and eq. (27) are equivalent boils down to showing that *e^kτ_D_^* = (*α* + *k*)/*α*. We call *τ_B_* the time spent by a bacterium in stage B. In the CH model the times *τ_B_*, *τ_C_* are constant. The division time is *τ*_div_ = *τ_B_* + *τ_C_* + *τ_D_*. The population growth rate is *k* = ln2/*τ*_div_.

Since the division time is constant, the steady-state age distribution is expressed by

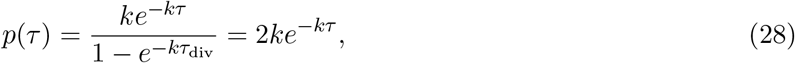

see e.g. Powell (1956)

**Figure B1:**
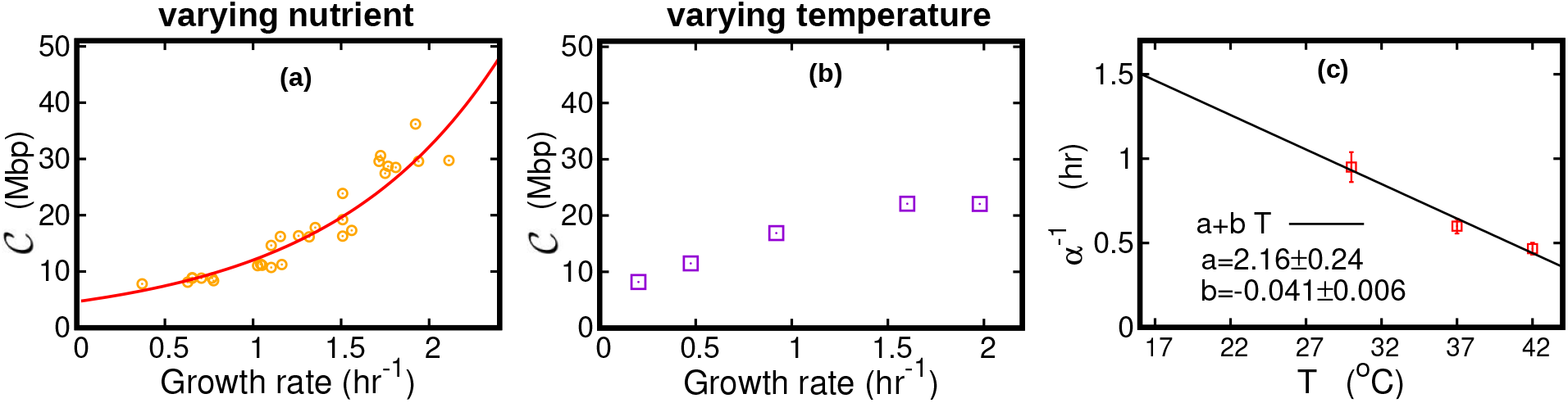
DNA content per cell as function of the growth rate for the case of varying nutrient in (a) and of varying temperature in (b). In (a), the experimental data (orangle circles) are from (Si et al., 2017). The solid red line is from (27), in which we used *τ_C_* = 38 min and *τ_D_* = 37.1 min (Si et al., 2017). The curve in (b) is from (26). In this case we substituted velocity of replisomes and growth rate of cells at each temperature in (26). In addition, we assumed a linear temperature dependence for the post replication duration (*α*^−1^), see (c). The parameters of the linear fit are determined from the data (red squares) reported in Stokke et al. (2012) for the LuriaBroth medium. We used this linear fit to extrapolate the value of **α** for different temperatures in (26).

We now express the steady-state fractions *f_B_*, *f_C_*, *f_D_* of cells in stage B, C, D as

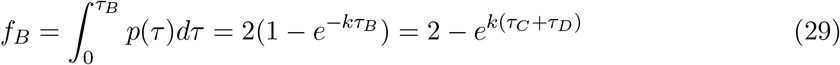

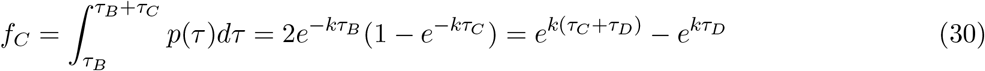

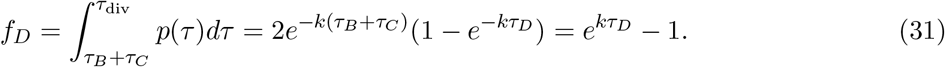

The ratio of the number of post replication genomes to the number of templates is equal to the fraction of cells in stage D. Therefore, equating the fraction *f_D_* with *N_P_/N_T_* = *k*/*α* obtained from eqs. (7) and (9) of the Main Text, we obtain *e^kτ_D_^* = (*α* + *k*)/*α* as expected.

## Appendix C Comparison between the average speed estimated from the oscillatory speed model and constant speed model

**Table C1:** Comparison between average (over sample to sample variation) speed estimated in the constant and oscillatory speed models. Temperature are expressed in Celsius and speeds in bp s^−1^. Last column shows the average growth rate (expressed in h*r*^−1^) at different temperatures.

| Temperature | $\bar{v}$ (constant speed) | $\bar{v}$ (oscillatory speed) | $k$ (growth rate) |
| --- | --- | --- | --- |
| 17 | $221 \pm 17$ | $243 \pm 36$ | $0.20 \pm 0.02$ |
| 22 | $373 \pm 29$ | $350 \pm 28$ | $0.47 \pm 0.04$ |
| 27 | $528 \pm 31$ | $542 \pm 30$ | $0.92 \pm 0.05$ |
| 32 | $812 \pm 64$ | $823 \pm 65$ | $1.60 \pm 0.12$ |
| 37 | $961 \pm 51$ | $972 \pm 51$ | $1.98 \pm 0.10$ |

## Appendix D Estimate of the diffusion coefficient from the Stokes-Einstein relation

The Stokes-Einstein relation expresses the diffusion coefficient of a spherical particle immersed in a fluid (Hynes, 1977). If *r* is the radius of the particle, *η* and *T* are viscosity and the temperature of the fluid respectively, then according to the Stokes-Einstein relation, the diffusion coefficient is

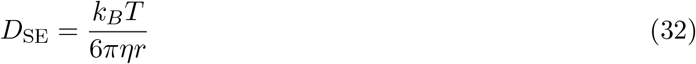

where *k_B_* is the Boltzmann constant. The radius of an E. Coli replisome is *r* ≈ 50 *nm* (Reyes-Lamothe et al., 2010), viscosity of water is *η* = 0.7mPa s at room temperature *T* = 310K. Using that *k_B_* = 1.38 × 10^−23^*JK*^−1^, we estimate a diffusion coefficient of replisomes in water equal to *D*_SE,W_ = 6*μ*m^2^s^−1^. The typical base pair distance is 3.4*A*°. Therefore in terms of base-pair (bp), *D*_SE,W_ ≈ 56kbp^2^s^−1^. The diffusion constant of large macromolecules in the cytoplasm is found to be about 10 times smaller than in water (Verkman, 2002). This results in an estimate of the diffusion constant of replisomes in the cytoplasm of *D*_SE,C_ ≈ 6kbp^2^s^−1^, as reported in the Main Text.

## Appendix E Exact solution in the oscillatory velocity case for *D* = 0

In this Section we present the exact solution to the oscillatory velocity model in the absence of diffusion (*D* = 0). For an arbitrary choice of the function *v*(*x*) and *D* = 0, the steady state solution of eq. (11) in the Main Text reads

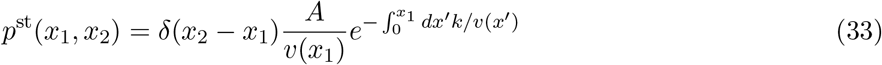

where *A* is a normalization constant that ensures 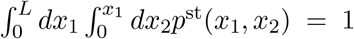. The rate at which replication completes is 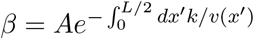. From eq. (14) of the Main Text, we obtain

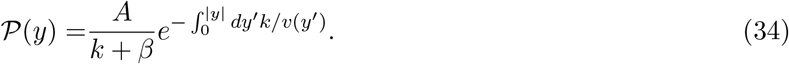

For the specific form of *v*(*x*) given in eq. (3) in the Main Text, the integral in (34) is equal to

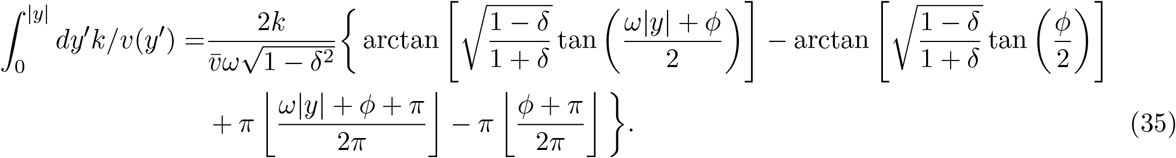

where ⌊·⌋ is the floor function. We use the expression of this integral in (34) and substitute the result in eq. (15) of the Main Text to obtain the DNA abundance distribution. We computed the normalization factor of the DNA abundance distribution (i.e., the denominator of eq. (15) of the Main Text) by numerical integration.

## Supporting Figures

**Figure S1:**
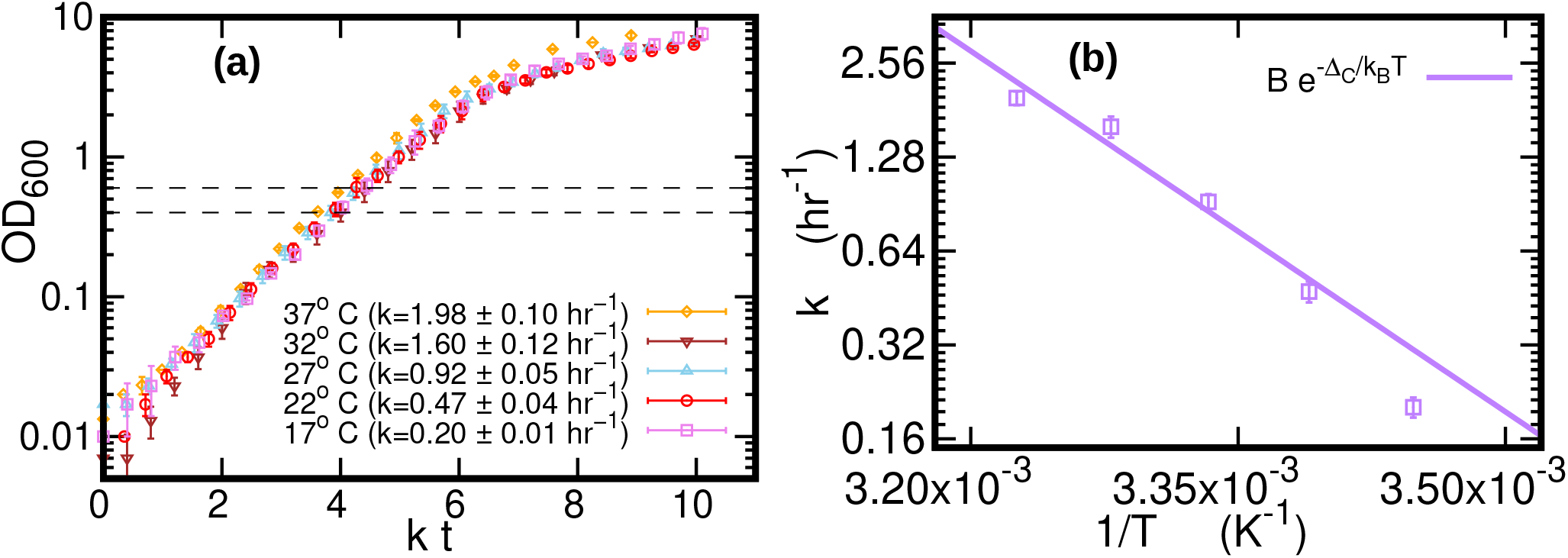
(a) Optical density as a function of time, scaled by the growth rate *k*. The data is averaged over three different replicates at each temperature. Error bars represent standard deviations. Dashed lines mark the OD window in which the cells are harvested. (b) Average growth rate as a function of temperature. The growth rate *k_i_* for each sample *i* = 1, 2, 3 at a given temperature is computed by fitting the optical density to a logistic function *a_i_*/[1 + *b_i_* exp(–*k_i_t*)], where *a_i_* and *b_i_* are sample-specific constants (Zwietering et al., 1990). The growth rate *k* is the average of the *k_i_*s. The solid purple line is an Arrhenius fit to the data, resulting in *B* = (6.0 ± 24.9) × 10^12^hr^−1^ and Δ_*C*_ = (74 ± 10)kJmol^−1^. We exclude the data point for *T* = 17°C from the fit.

**Figure S2:**
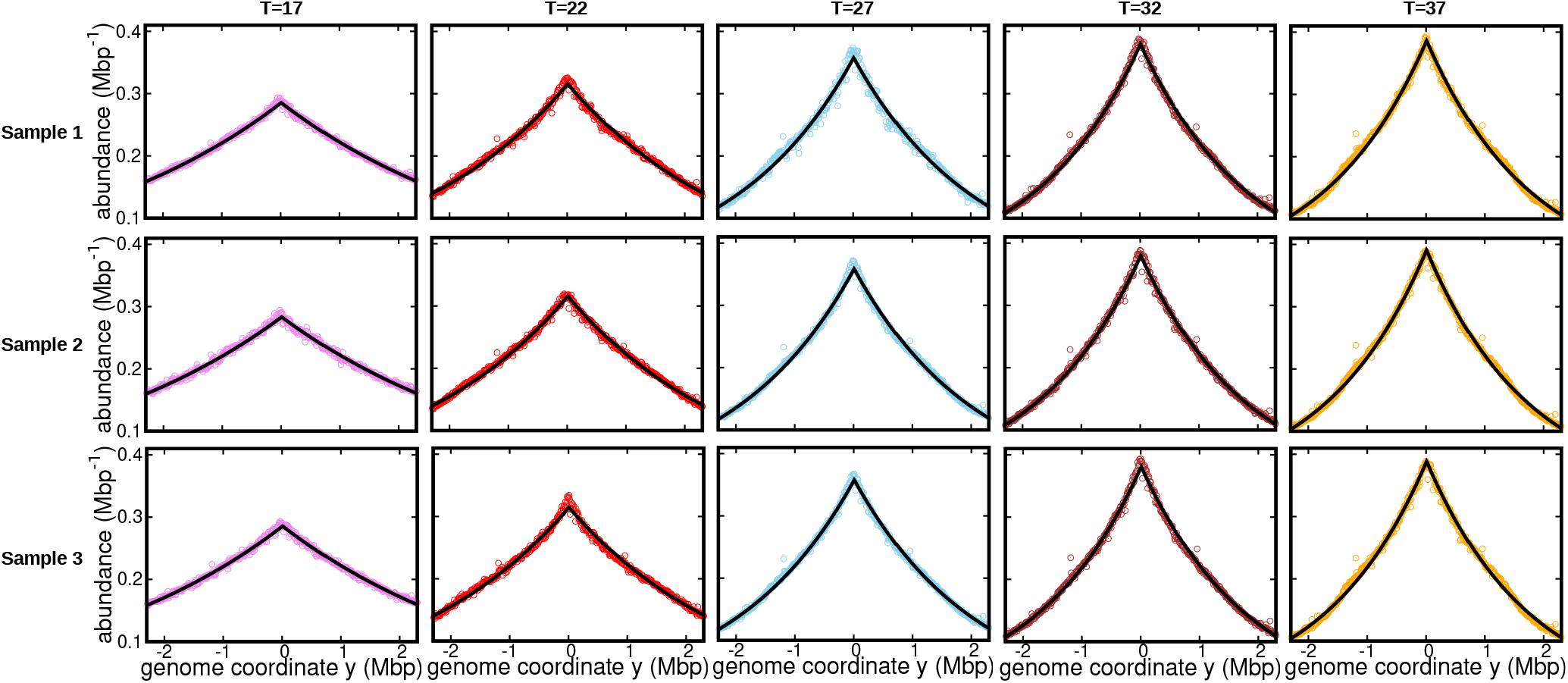
DNA abundance distribution at different temperatures. Bias elimination is carried out using the DNA abundance of the stationary cells grown at temperature 17°*C*. Symbols are from experiments and the solid black line is predictions of the constant velocity model. The DNA abundance distribution presented in Figure 1d, Figure 2a, and Figure 3 of the Main Text corresponds to Sample 1 (top panel).

**Figure S3:**
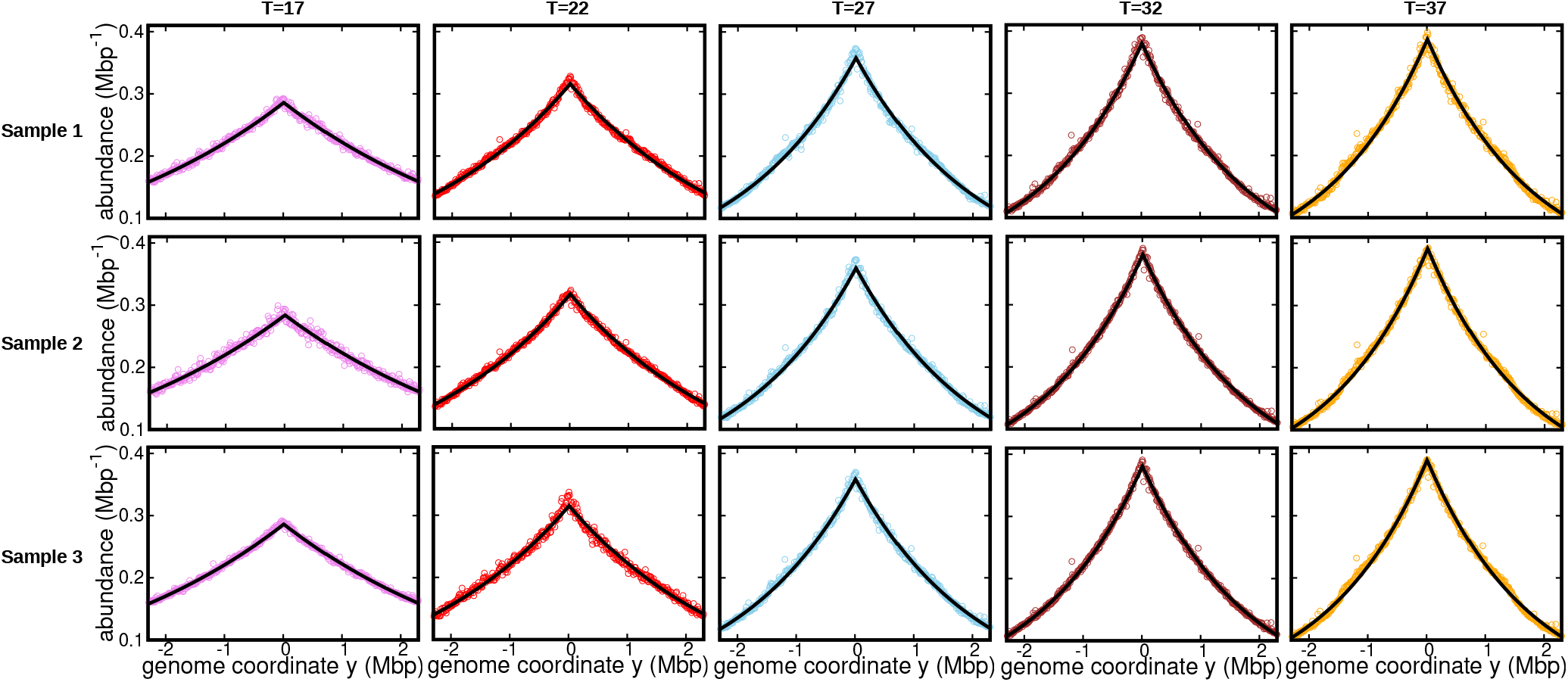
DNA abundance distribution at different temperature. Bias elimination is carried out using the DNA abundance of the stationary cells grown at temperature 27°*C*. Symbols are from experiments and the solid black line is predictions of the constant velocity model.

**Figure S4:**
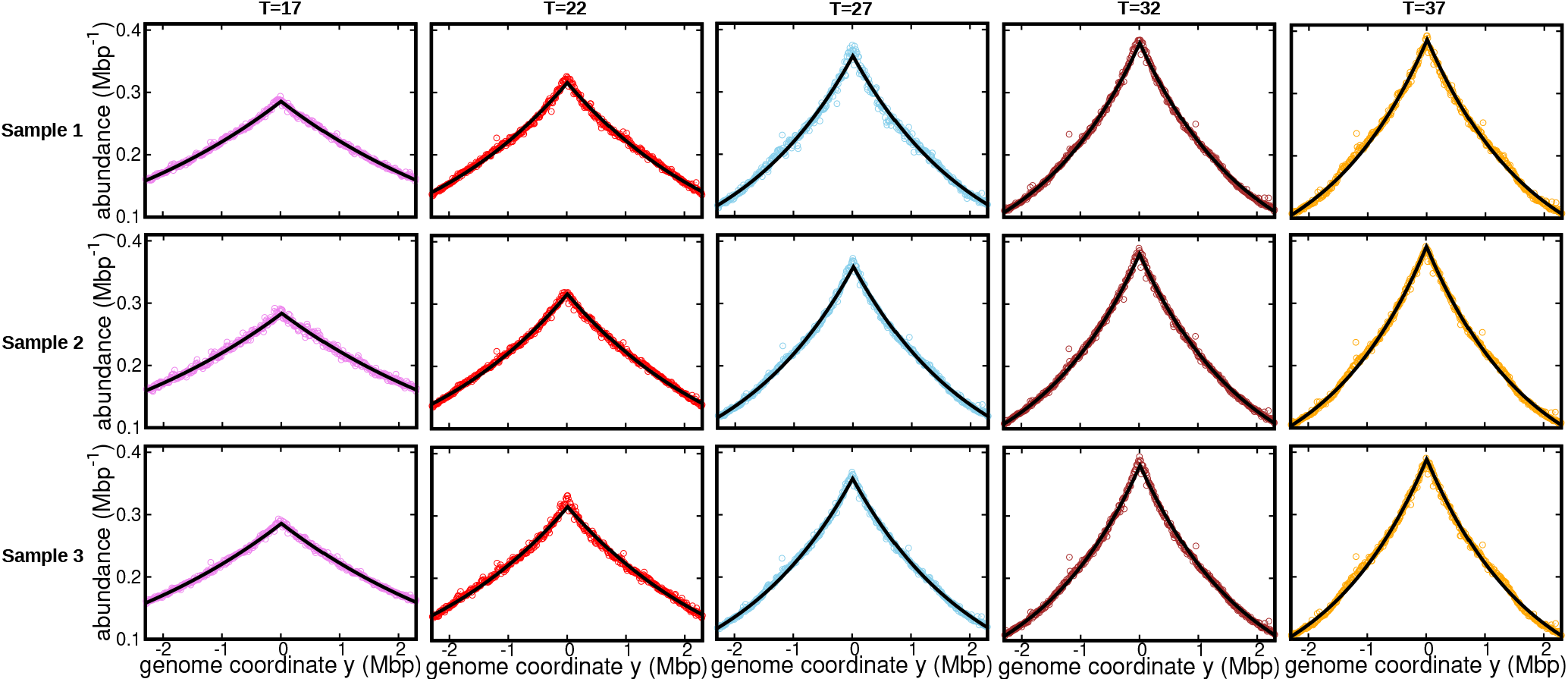
DNA abundance distribution at different temperature. Bias elimination is carried out using the DNA abundance of the stationary cells grown at temperature 37°*C*. Symbols are from experiments and the solid black line is predictions of the constant velocity model.

**Figure S5:**
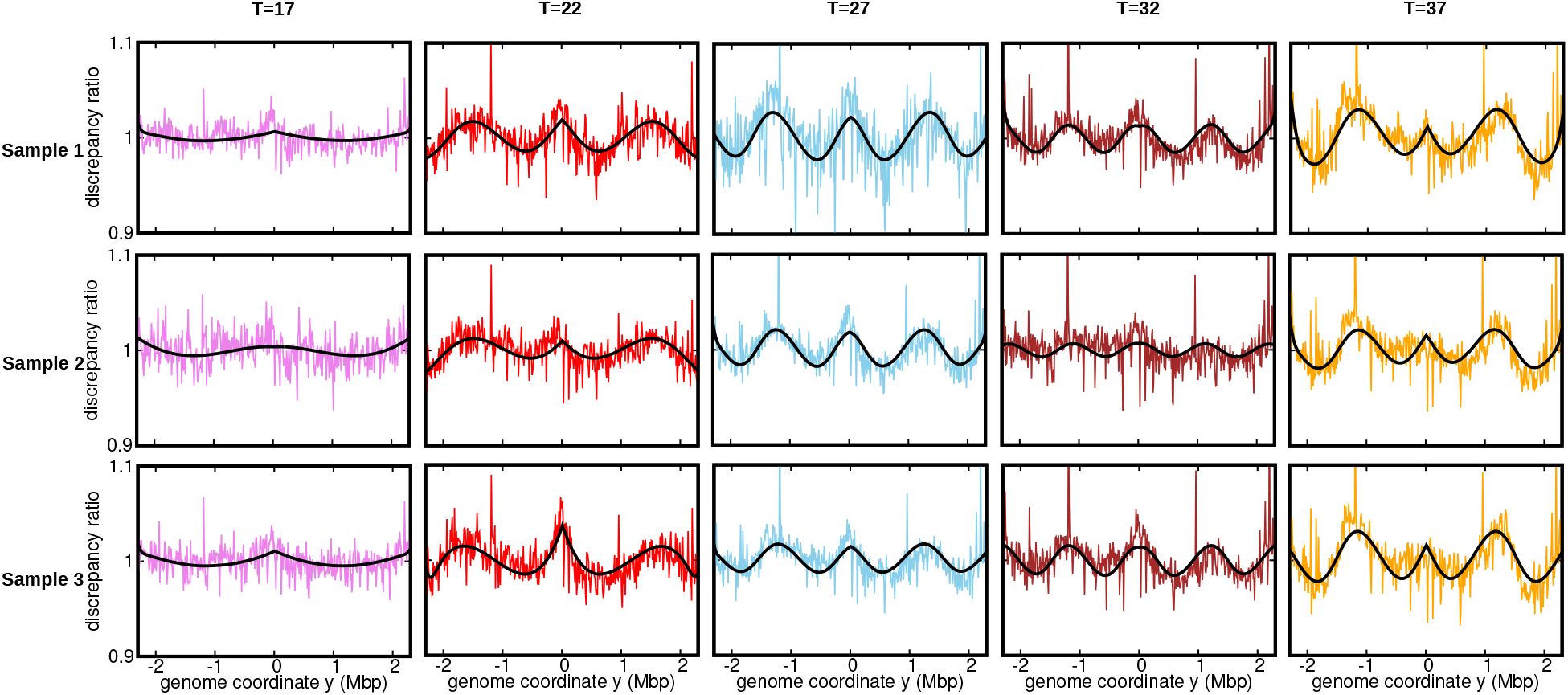
Ratios of the experimental DNA abundance over the corresponding prediction assuming constant speed and *D* = 0. The solid black lines represent the ratios of the predictions assuming oscillatory speed over constant speed. The data here is for the stationary temperature 17°*C*. The discrepancy ratios presented in Figure 3 of the Main Text is for Sample 1 (top most panel).

**Figure S6:**
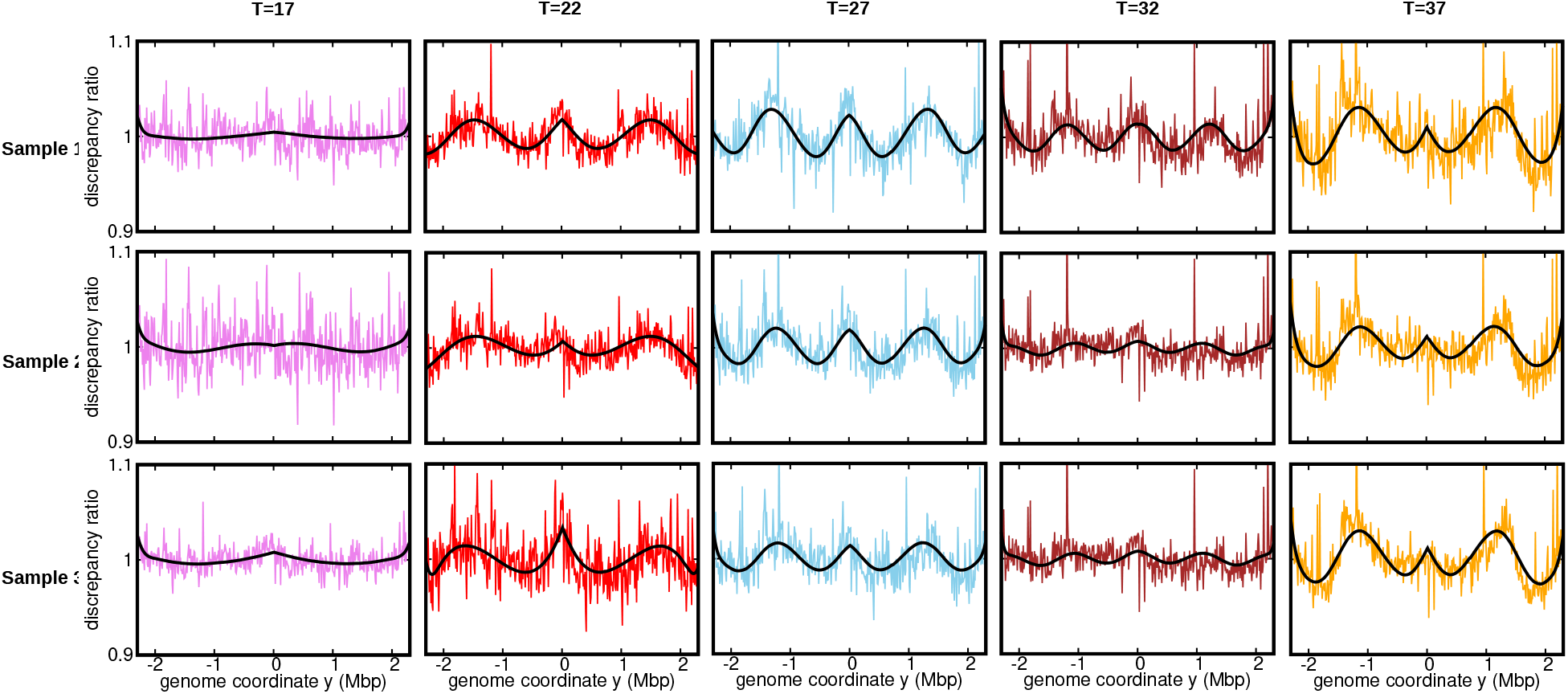
Ratios of the experimental DNA abundance over the corresponding prediction assuming constant speed and *D* = 0. The solid black lines represent the ratios of the predictions assuming oscillatory speed over constant speed. The data here is for the stationary temperature 27°*C*.

**Figure S7:**
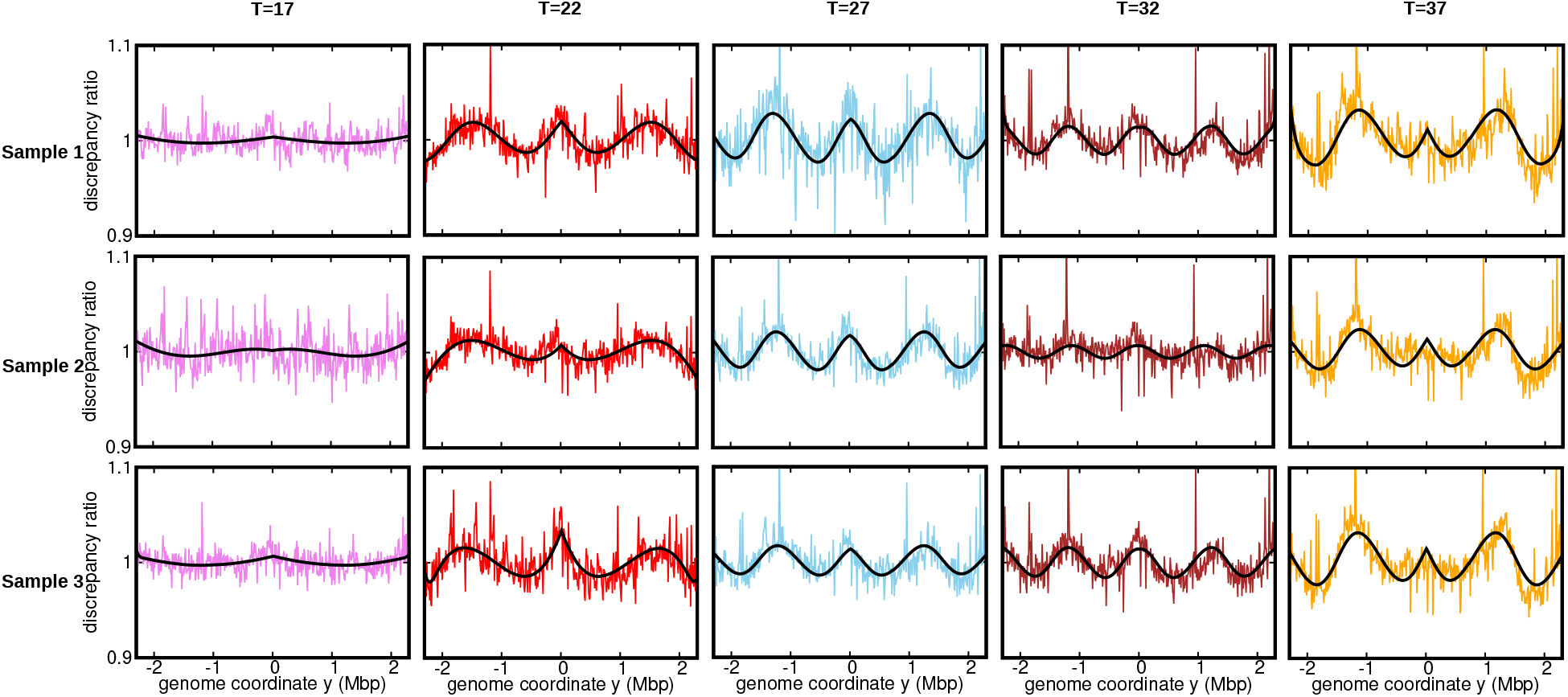
Ratios of the experimental DNA abundance over the corresponding prediction assuming constant speed and *D* = 0. The solid black lines represent the ratios of the predictions assuming oscillatory speed over constant speed. The data here is for the stationary temperature 37°*C*.

**Figure S8:**
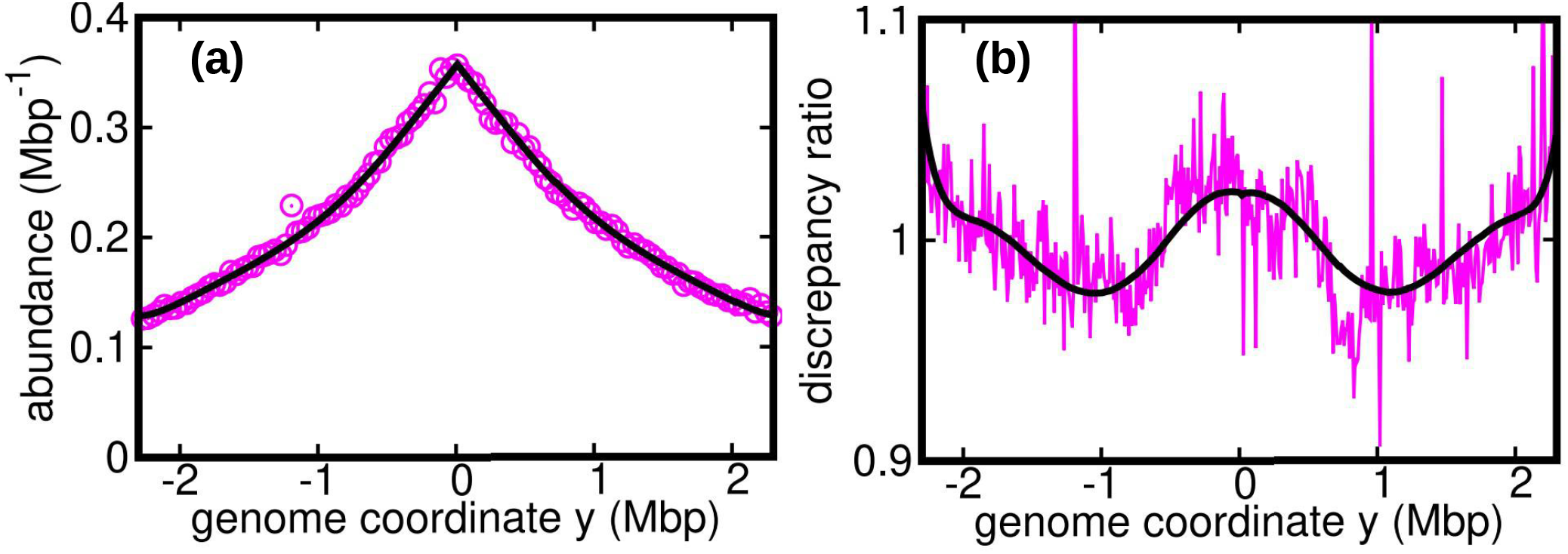
(a) DNA abundance distribution from the data reported in Midgley-Smith et al. (2018b) for *E. Coli* growing in a Luria broth medium. The solid black line is the prediction from the constant speed model. The reported growth rate for this culture is *k* = 2.15 ± 0.18hr^−1^ (Midgley-Smith et al., 2018b). Substituting this value in our model we obtain the average speed *v* = 1306 ± 115bps^−1^. (b) Ratios of the experimental DNA abundance over the prediction of the constant speed model. The solid black line represents the ratios of the predictions assuming oscillatory speed over constant speed. The estimated parameters of the oscillatory velocity model are 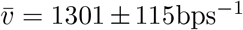, *δ* = 0.14 ± 0.02, *ω* = 3.2rad Mbp^−1^, *ϕ* = 1.3rad and *D* = 16 ± 14kbp^2^s^−1^.

**Figure S9:**
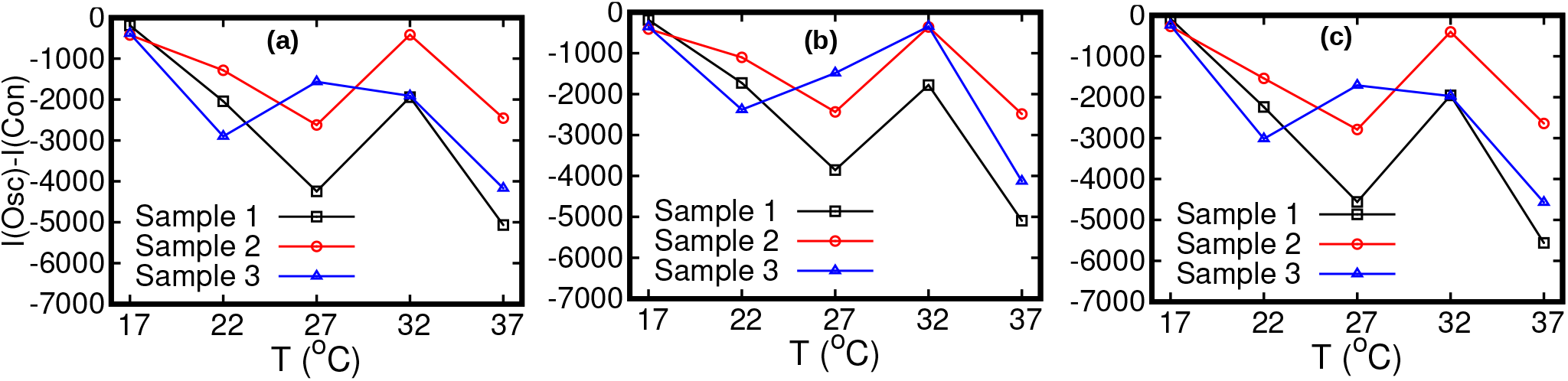
Statistical comparison between the constant velocity model and oscillatory model by means of the Akaike information criterion. The Akaike information is defined as 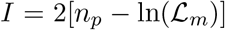, where *n_p_* is the number of fitted parameters in a model and 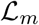 is the maximum likelihood. The constant velocity model includes a single free parameters (*n_p_* = 1) versus five (*n_p_* = 5) for the oscillatory velocity model. The Akaike information criterion prescribes that, given a set of candidate models, one should prefer the one characterized by the minimum Akaike information. The Akaike information for the constant velocity model, *I*(Con), is larger than that for the the oscillatory velocity model, *I*(Osc), for all temperatures. The stationary data used for bias elimination correspond to temperatures (a) 17°C, (b) 27°C, and (c) 37°C.

**Figure S10:**
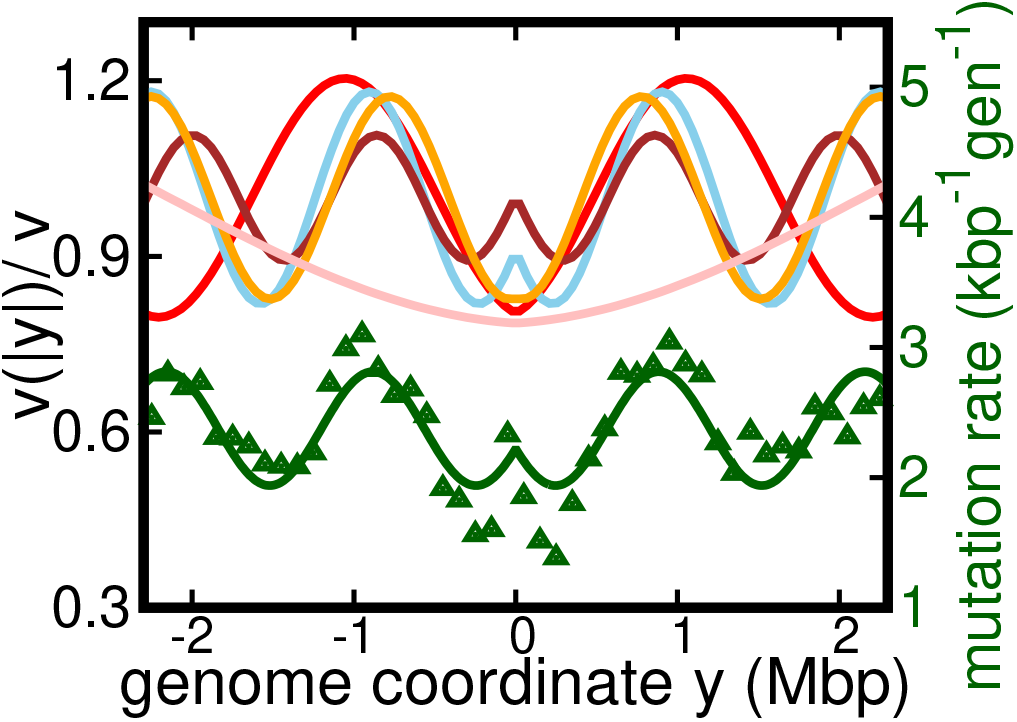
Results of the oscillatory speed model. This figure is same as the Figure 4a of the Main Text, but with additional curve for *T* = 17°C. Solid lines: relative speeds 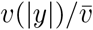 along the genome (Pink: *T* = 17°C, Red: *T* = 22°C, sky blue: *T* = 27°C, brown: *T* = 32°C, and orange: *T* = 37°C).

**Figure S11:**
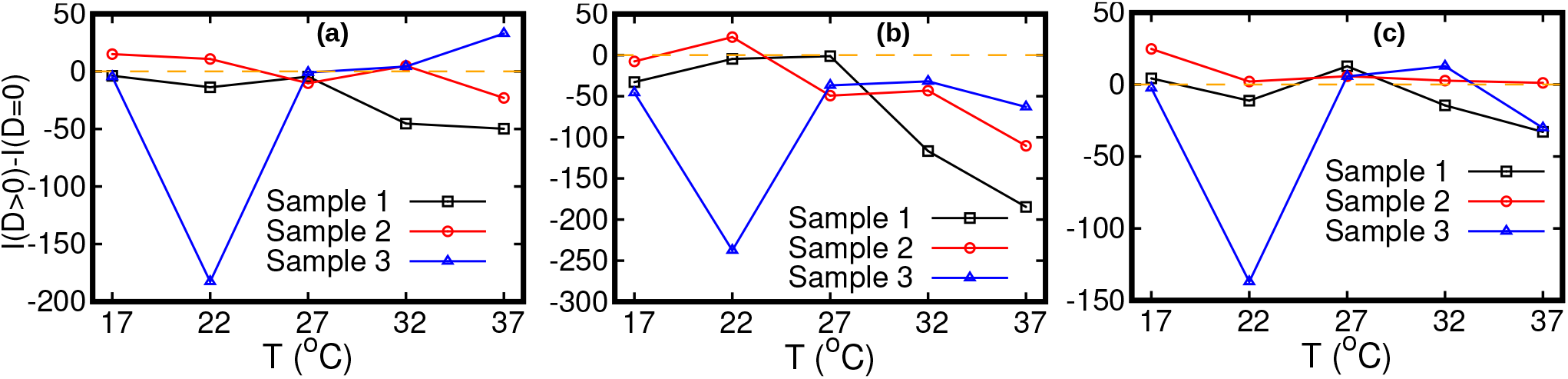
Statistical comparison between the oscillatory model with diffusion (*D* > 0) and without diffusion (*D* = 0) by means of the Akaike information criterion. The number of parameters is four (*n_p_* = 4) for *D* = 0 and five (*n_p_* = 5) for *D* > 0. Difference between Akaike information *I* for *D* > 0 and *D* = 0. If this difference is negative, the model with *D* > 0 should be preferred. The stationary data used for bias elimination correspond to temperatures 17°C in (a), 27°C in (b) and 37°C in (c).

